# Optimized Quantitative Susceptibility Mapping at 7T MRI for Assessing Iron deposition in Alzheimer’s Disease

**DOI:** 10.1101/2025.08.31.673334

**Authors:** Felisha Ma, Pinar Senay Özbay, Berkin Bilgic, Trey Hedden, Bradley Delman, Priti Balchandani, Akbar Alipour

**Affiliations:** BioMedical Engineering and Imaging Institute (BMEII), Icahn School of Medicine at Mount Sinai, New York, NY 10029, US; Institute of Biomedical Engineering, Bogazici University, Istanbul, Turkiye; Athinoula A. Martinos Center for Biomedical Imaging, Massachusetts General Hospital and Harvard Medical School, Boston, MA 02129, US; Department of Neurology, Icahn School of Medicine at Mount Sinai, New York, NY 10029, US; Department of Diagnostic, Molecular and Interventional Radiology, Icahn School of Medicine at Mount Sinai, New York, NY 10029, US

## Abstract

**INTRODUCTION:** Elevated brain iron levels are common in Alzheimer’s disease (AD). Quantitative Susceptibility Mapping (QSM) is an advanced MRI technique for assessing iron accumulation. The optimized QSM at 7 Tesla (7T) MRI may further improve the sensitivity to detect subtle susceptibility changes in AD.

**METHODS:** We optimized a QSM processing pipeline for 7T MRI by systematically comparing multiple reconstruction algorithms. Evaluation criteria included image quality, artifact suppression, and anatomical clarity. The finalized pipeline was applied to individuals with AD and healthy controls (HCs).

**RESULTS:** The results revealed significantly elevated magnetic susceptibility values in the globus pallidus and dentate nucleus of the AD group compared to HCs. These findings were confirmed through both visual inspection and quantitative analysis of high-resolution QSM maps.

**DISCUSSION:** Our results highlight the importance of optimizing QSM pipelines at 7T for accurate susceptibility quantification. We identified an optimal pipeline suitable for future applications in patients with AD and other neurological conditions.

## INTRODUCTION

Alzheimer’s disease (AD) is an age-related neurodegenerative disorder and the leading cause of dementia in older adults. It is characterized by progressive cognitive decline that significantly impacts memory, thinking, and behavior [1, 2]. Two of the main pathological hallmarks of AD are the accumulation of amyloid-beta (Aβ) plaques and the formation of tau protein tangles in the brain [3]. These microscopic changes are believed to begin years before noticeable neurological symptoms appear such as memory loss, disorientation, or difficulty with language, preceding gross morphological changes like hippocampal atrophy [4, 5]. Currently available biomarkers including positron emission tomography (PET) imaging, cerebrospinal fluid (CSF), and blood plasma enable the detection of Aβ and tau pathology for the diagnosis and staging of AD [6-9]. However, these methods have limited sensitivity in the earliest stages of disease [10]. Previous studies have shown that iron deposition is elevated in the brains of individuals with AD and may play a critical role in the disease’s pathogenesis [11-15]. Furthermore, iron has been found to interact with hallmark AD proteins, promoting Aβ aggregation and tau hyperphosphorylation, both of which are central to AD pathology [16-20]. These findings suggest that abnormal brain iron levels are not merely a consequence of neurodegeneration but may actively drive disease progression. Neuroimaging MRI techniques, such as T2*-weighted imaging, R2* mapping, and susceptibility-weighted imaging (SWI), have enabled the in vivo evaluation of brain iron level, providing insights into its role in neurodegeneration [11, 14, 21]. T2* and R2* only offer indirect and non-specific signal decay rates affected by various susceptibility sources, including iron, calcification, and myeline. SWI is highly sensitive but lacks quantitative capability and may be influenced by veins or hemorrhage and blooming artifacts, leading to false interpretations.

Quantitative Susceptibility Mapping (QSM) is an advanced MRI technique that enables the in vivo measurement of tissue iron levels by detecting magnetic susceptibility changes associated with iron deposition in brain tissue [22, 23]. QSM applications span a wide range of clinical and research domains, including Parkinson’s diseases, multiple sclerosis, stroke, brain tumors, and cardiac disease, where it aids in assessing iron overload and myocardial tissue characterization [24-28]. QSM holds significant potential for detecting and monitoring iron accumulation in AD, offering a non-invasive, in vivo method to quantify and localize iron deposits in affected brain regions such as the hippocampus, basal ganglia, thalamus, substantia nigra, and cortical areas, particularly within the temporal and frontal lobes [29-32].

Ultra-high-field (UHF) MRI at 7 Tesla (7T) significantly enhances imaging capabilities by providing a higher signal-to-noise ratio (SNR) and improved contrast compared to conventional 1.5T and 3T systems [33-35]. This increased SNR enables sub-millimeter resolution imaging, offering greater sensitivity and more detailed visualization of cortical and deep brain structures. These advantages make 7T MRI particularly valuable for detecting subtle neuroanatomical changes that may be missed at lower field strengths. Additionally, the enhanced resolution and susceptibility sensitivity of 7T QSM make it especially well-suited for assessing iron accumulation in the brains of patients with AD [36-42].

QSM is performed as a series of post-processing steps including phase unwrapping, background field removal, and dipole inversion applied to MRI phase and magnitude images [23]. Previous studies have reported the use of QSM in AD at both 3T and 7T MRI, demonstrating its potential for detecting iron-related changes in the brain [21, 29, 31, 41-43]. Despite the progress made in standardizing QSM practices, there remains a clear need for the development of an optimized QSM pipeline specifically tailored for 7T MRI data. Artifacts such as increased phase wrapping, B_0_ inhomogeneities, susceptibility-induced distortions, and geometric distortion are more prominent at 7T MRI requiring adjustment strategies in each step [40, 42]. Therefore, a dedicated, standardized QSM pipeline optimized for 7T MRI is essential to fully harness the potential of 7T QSM in detecting subtle iron-related pathologies, improving the accuracy and clinical relevance in AD and other neurodegenerative disorders.

In this study, our primary goals were to (1) establish an optimized end-to-end reconstruction pipeline for 7T QSM in the brain, and (2) leverage optimized and enhanced QSM at 7T to study differences in brain iron aggregation between individuals with AD and healthy controls (HCs). Given that QSM is highly sensitive to factors such as magnetic field strength, spatial resolution, and acquisition parameters, careful protocol optimization at 7T is essential. This not only maximizes image quality and measurement accuracy but also represents a critical step toward ensuring reproducibility and standardization in UHF neuroimaging studies. By using rigorously optimized QSM at 7T, we aim to obtain more reliable and biologically meaningful assessments of regional iron accumulation associated with AD pathology.

Currently, the in vivo quantification of iron concentration in targeted brain regions represents a promising biomarker for advancing our understanding of the underlying mechanisms of the disease. As a result, brain iron is being explored as a potential biomarker for early diagnosis and a target for therapeutic intervention.

## METHODS

### Participants

Data from five healthy young adults (aged 30 ± 5 years) were used to optimize the imaging protocol and QSM processing pipeline prior to applying the methodology to the main study cohort. The primary study included 10 individuals with AD (ages 60–85; 6 female) and 10 HCs (ages 60– 85; 5 female). All participants were enrolled through the Mount Sinai Alzheimer’s Disease Research Center (ADRC), a longitudinal cohort study involving individuals identified through clinical practice. The study protocol was approved by the Institutional Review Board of the Icahn School of Medicine at Mount Sinai, and written informed consent was obtained from all participants in accordance with the Declaration of Helsinki. Each participant underwent a comprehensive clinical evaluation, including review of medical history, mental status examination, neurological examination, and neuropsychological testing. Clinical diagnoses of mild cognitive impairment and dementia were determined through expert panel review using established diagnostic criteria.

### Data Acquisition

MRI scanning was conducted on a 7T MRI (Magnetom, Siemens Healthineers, Erlangen, Germany) system using a 1Tx/32Rx Nova head coil. T1-weighted (T1w) imaging was performed using a 3D Magnetization Prepared Two Rapid Acquisition Gradient Echo (MP2RAGE) sequence with the following parameters: repetition time (TR) = 6000 ms, echo time (TE) = 3.62 ms, inversion time (TI_1_/TI_2_) = 1050/3000 ms, matrix size = 256 × 146, flip angles = 5° and 4° for the first and second inversion times, respectively, and an isotropic spatial resolution of 0.7 × 0.7 × 0.7 mm^3^. For QSM analysis, the 3D multi-echo gradient echo (3D-MEGRE) sequence was used, consisting of six echoes with the following acquisition parameters: TR = 28 ms; TE = 4, 8, 12, 16, 20, and 24 ms; flip angle = 12°; and GRAPPA factor = 2. To assess the effect of spatial resolution on QSM reconstruction, scans were performed at five different resolutions: 0.2 × 0.2 × 1.5 mm^3^, 0.3 × 0.3 × 1.5 mm^3^, 0.5 × 0.5 × 1.5 mm^3^, 0.7 × 0.7 × 0.7 mm^3^, and 0.7 × 0.7 × 1.5 mm^3^ with the acquisition time of 10.1 minutes, 8.4 minutes, 8.1 minutes, 7.9 minutes, and 7 minutes. These varying resolutions were used to evaluate the trade-off between image quality and acquisition efficiency for optimal QSM pipeline development. The optimized protocol and reconstruction pipeline were subsequently applied to a separate cohort consisting of individuals with AD and age-matched HCs to investigate the iron-related susceptibility differences between groups.

### Optimizing QSM Processing Pipeline

From a wide catalog of available algorithms, researchers can customize each step of QSM to suit specific study purposes. Previous studies have established that choice of different technique in QSM computational steps, particularly in phase-unwrapping, background field removal, and dipole inversion, as well as their parameters can differentially impact the final QSM output [44, 45]. Because QSM results are contingent upon all steps performing in tandem, with outcomes from intermediate stages influencing downstream results understanding how algorithms operate collectively becomes crucial. As a result, there is a need to investigate end-to-end pipelines for neuroimaging implementation at 7T. The QSM Consensus Organization Committee, under the International Society for Magnetic Resonance in Medicine (ISMRM) Electro-Magnetic Tissue Properties Study Group, has published a comprehensive set of recommendations for implementing QSM in clinical brain research [46]. This consensus aims to standardize QSM methodologies to ensure repeatability, reproducibility, and reliability across studies. In this work, we followed these guidelines to optimize both the data acquisition and post-processing pipelines, with the goal of facilitating the robust and consistent use of QSM in UHF systems.

We utilized an open-source processing pipeline tool called SEPIA, which is specifically designed for the post-processing of MRI phase images and QSM. SEPIA integrates multiple publicly available QSM toolboxes, offering enhanced flexibility and modularity in QSM processing [44]. QSM reconstruction was performed in Matlab R2022b (Mathworks, Natick, USA) using the SEPIA toolbox (v1.2.2.4) on a macOS (2.3 GHz 8-Core Intel Core i9 CPU, 32GB RAM). SEPIA offers multiple phase unwrapping algorithms, background field removal methods, and dipole inversion techniques available for QSM reconstruction. In this study, overall, 32 reconstruction pipelines were generated for comparison from combinations of two phase-unwrapping, four background field removal, and four dipole inversion algorithms. Figure 1 shows the schematic of the QSM processing workflow that we used. The unwrapped phase data, along with the brain mask, were used to perform background field removal isolating the local tissue-induced magnetic field variations. A brain mask was generated for effective background field removal using magnitude data. Finally, susceptibility maps were computed by applying an inversion algorithm to solve the dipole deconvolution problem.

**Figure 1.**
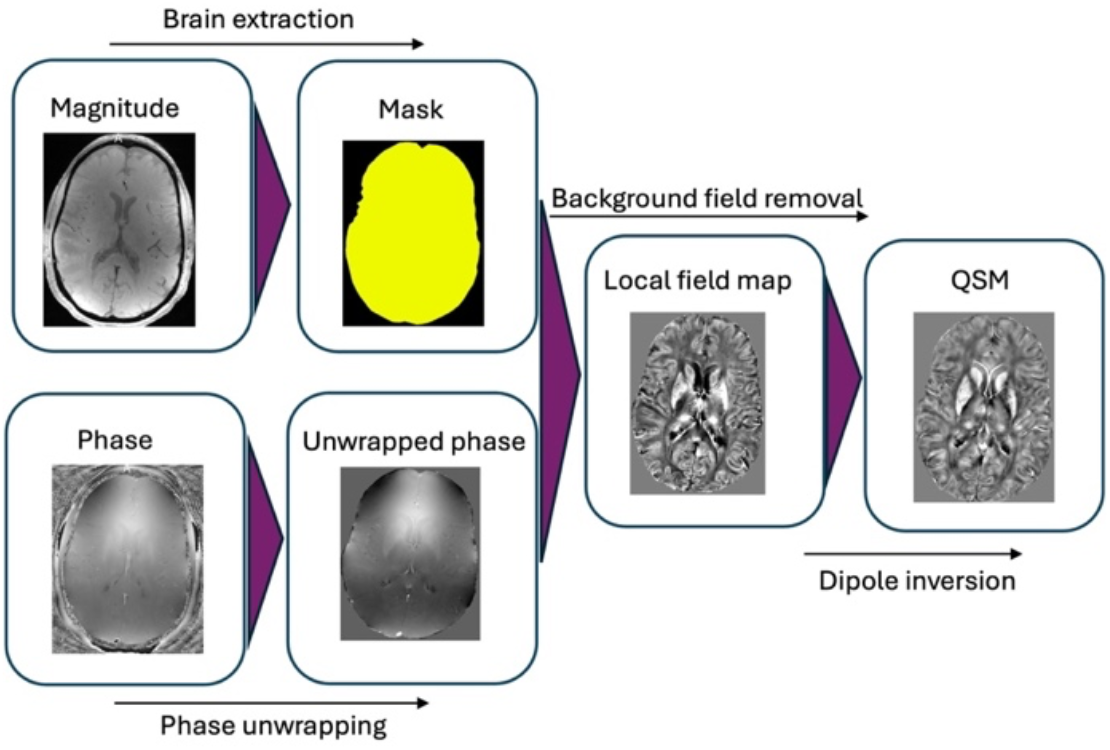
QSM processing pipeline. A brain mask from magnitude images and unwrapped phase data were used for background field removal, followed by dipole inversion to generate susceptibility maps for quantitative analysis.

For QSM processing, we implemented an exact phase unwrapping method to ensure high-fidelity reconstruction of the local magnetic field. Consistent with recommended best practices, we performed echo combination prior to background field removal. To enhance SNR, we used weighted echo averaging following template-based phase unwrapping and explicit phase offset removal. This strategy was selected based on its demonstrated robustness across different acquisition protocols and its effectiveness in producing reliable susceptibility maps, particularly in UHF imaging contexts. As phase unwrapping technique, the Laplacian method and the Speedy rEgion-Growing algorithm for Unwrapping Estimated phase (SEGUE) were employed, using optimal echo phase combination [47-49].

We utilized the Brain Extraction Tool (BET) from the FMRIB Software Library (FSL) to perform brain masking [50]. Although often underreported, masking is a critical step in the QSM pipeline, particularly for effective background field removal. By accurately defining the region of interest (ROI) within the field of view, masking ensures that field map computations are constrained to relevant anatomical structures. The BET algorithm provided an efficient and reliable approach for isolating brain tissue, thereby supporting robust and accurate background field removal and downstream susceptibility mapping. This step was essential for maintaining the integrity and consistency of our QSM analysis. Given the known limitations of BET when applied to UHF MRI such as 7T, primarily due to fileds inhomogeneities and increased susceptibility effects, we optimized the fractional intensity threshold (f) parameter to improve mask accuracy. Specifically, we used a reduced value (f = 0.3), which provided better contouring of the brain boundaries in our 7T images.

As part of our QSM processing pipeline, we implemented background field removal to isolate the tissue field from the total field map. This step is essential because background fields, generated by susceptibility sources outside the brain, such as air that can be significantly stronger than the susceptibility differences within brain tissue at UHF. Background field removal was carried out using multiple techniques, including Variable-kernel Sophisticated Harmonic Artifact Reduction for Phase data (VSHARP) [48], Regularization-enabled Sophisticated Harmonic Artifact Reduction for Phase data (RESHARP) [51], Laplacian Boundary Value (LBV) [52], and Projection onto Dipole Field (PDF) [53]. Each method included detailed parameter settings, such as the size or radius of the Spherical Mean Value (SMV) kernel, tolerance levels, and polynomial fitting order. To avoid shadowing artifacts and improve the accuracy of dipole inversion, we selected a harmonic-based method that leverages the fact that background fields satisfy the Laplace equation within the brain. This approach enables effective separation of the local tissue-induced field from background contributions, thereby enhancing the reliability of the resulting susceptibility maps.

The final step in our QSM processing pipeline was dipole inversion, which involves spatial deconvolution of the tissue field using the dipole kernel to estimate the underlying tissue susceptibility distribution. This step is critical but inherently ill-posed due to the dipole kernel approaching zero near the magic angle (∼54.7°), which leads to numerical instability. To mitigate the impact of low SNR and deviations from ideal dipole behavior, common in certain brain regions, we applied masking to exclude problematic voxels, thereby reducing streaking and shadowing artifacts and improving the accuracy of the susceptibility maps. Dipole inversion was performed using several algorithms: Morphology Enabled Dipole Inversion (MEDI) [54], Iterative Least Squares (iLSQR) [48], Streaking Artifact Reduction for QSM (Star-QSM) [55], and Thresholded k-space Division (TKD) [56]. Parameters included regularization weights (lambda), reference tissue (typically CSF), zero-padding, edge mask thresholds, and optimization criteria. The algorithm and associated parameters are detailed in Table 1.

**Table 1:**
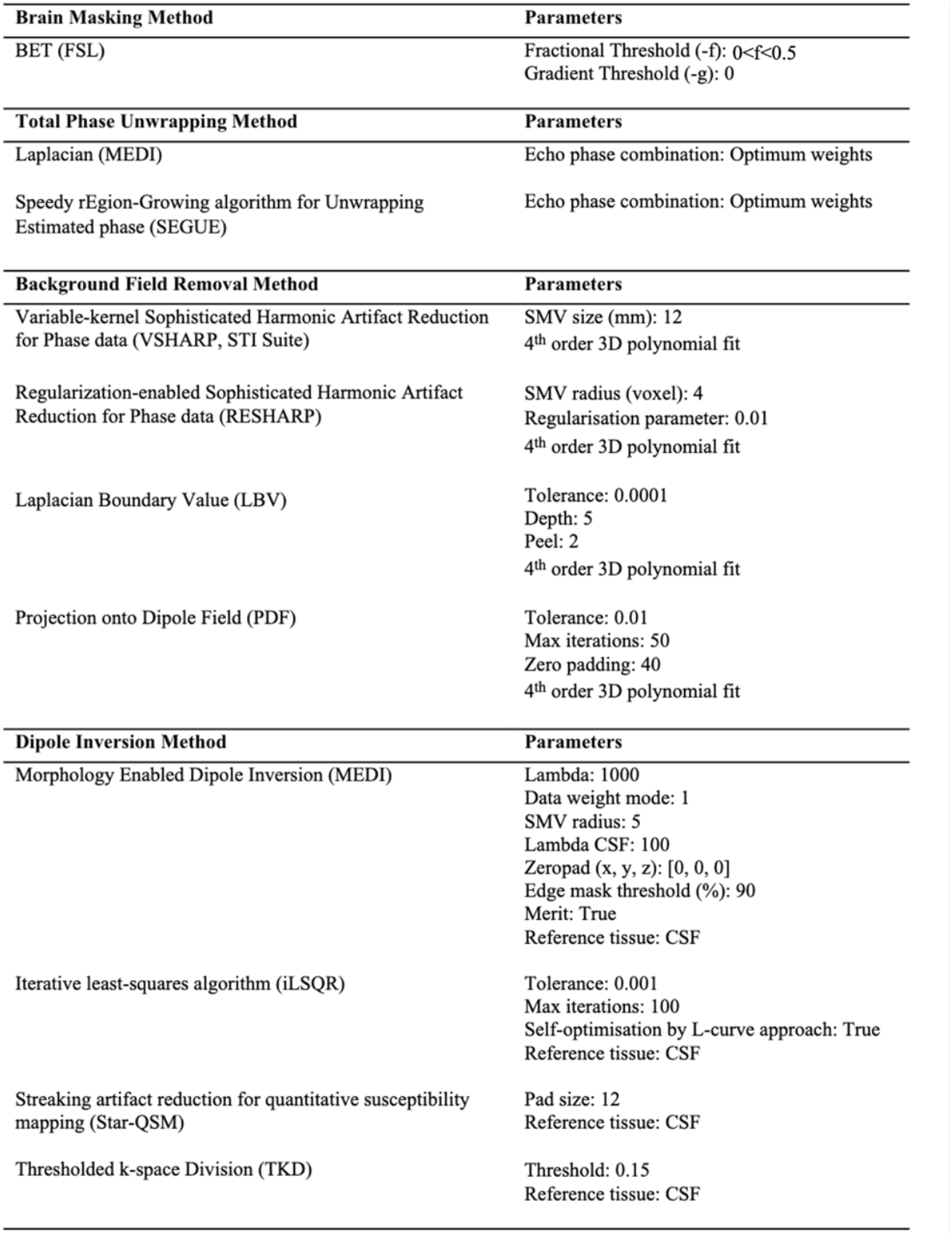
QSM reconstruction algorithms and parameters.

To further enhance uniformity and improve downstream processing, we applied bias field correction to the magnitude images prior to brain extraction. Bias field correction is a crucial pre-processing step, as it significantly impacts both qualitative and quantitative image analysis, particularly at high field strengths such as 7T. In this study, we used SPM’s bias field correction method, which helped mitigate the filed inhomogeneity at 7T MRI [57]. This correction improved the performance of brain masking and contributed to more consistent results in the subsequent QSM reconstruction.

To evaluate the impact of post-processing parameters on susceptibility quantification, we assessed two key factors: referencing strategy and voxel residual threshold (VRT). QSM maps were reconstructed using both CSF referencing and whole-brain mask referencing. CSF referencing involves setting the mean susceptibility of the CSF-typically assumed to be near zero-as the reference baseline, while brain mask referencing uses the mean susceptibility across the entire brain. In the context of SEPIA, the VRT is a parameter used during dipole inversion step to control the quality of the susceptibility map by managing noisy or unreliable voxels. By excluding these noisy or unreliable voxels, the VRT enhances the robustness and accuracy of the final susceptibility map, reducing artifacts and improving overall image quality. To examine the influence of brain masking criteria, we tested three different VRT values (0.3, 0.5, and 0.7) during the inversion process.

The mean magnetic susceptibility values and standard deviations (SD) were quantified from the QSM maps using the corresponding segmentation files generated by FreeSurfer software [58]. These values were then averaged across subjects for each ROI and referenced to the whole-brain mean susceptibility. The results were also evaluated through visual inspection with a focus on overall image quality. Given that both spatial resolution and the choice of reference region can influence image clarity and susceptibility estimation, we optimized acquisition parameters by testing five different resolutions and two reference tissue options on a preliminary sample set. Image quality assessment criteria included: (1) clarity of anatomical boundaries, which ensures accurate susceptibility quantification and reliable tissue differentiation; (2) presence of artifacts such as streaking, shading, or geometric distortion, which typically result from reconstruction errors and reduce interpretability; and (3) image uniformity, particularly relevant at 7T where magnetic field inhomogeneities can lead to uneven image intensities.

### Assessing the Brain Iron Level in AD

To assess brain iron levels in individuals with AD, we applied the optimized QSM processing pipeline to MRI data from 10 participants with AD and 10 age-matched HCs, all aged 65–80 years and meeting 7T MRI safety criteria. QSM was reconstructed from a 3D MEGRE sequence with a voxel size of 0.3 × 0.3 × 1.5 mm^3^, using the same imaging parameters as previously described. QSM quality and pipeline performance were evaluated based on anatomical boundary clarity, presence of artifacts, and image uniformity. All images were independently reviewed by two blinded neuroradiologists to ensure consistency and eliminate bias in qualitative assessment. Figure 2 shows axial susceptibility maps with six iron-rich brain regions manually segmented as ROIs using anatomical reference images. The selected ROIs included the putamen (PT), globus pallidus (GP), caudate nucleus (CN), red nucleus (RN), substantia nigra (SN), and dentate nucleus (DN). Susceptibility values were referenced to the mean susceptibility of CSF and compared between the AD and HC groups.

**Figure 2.**
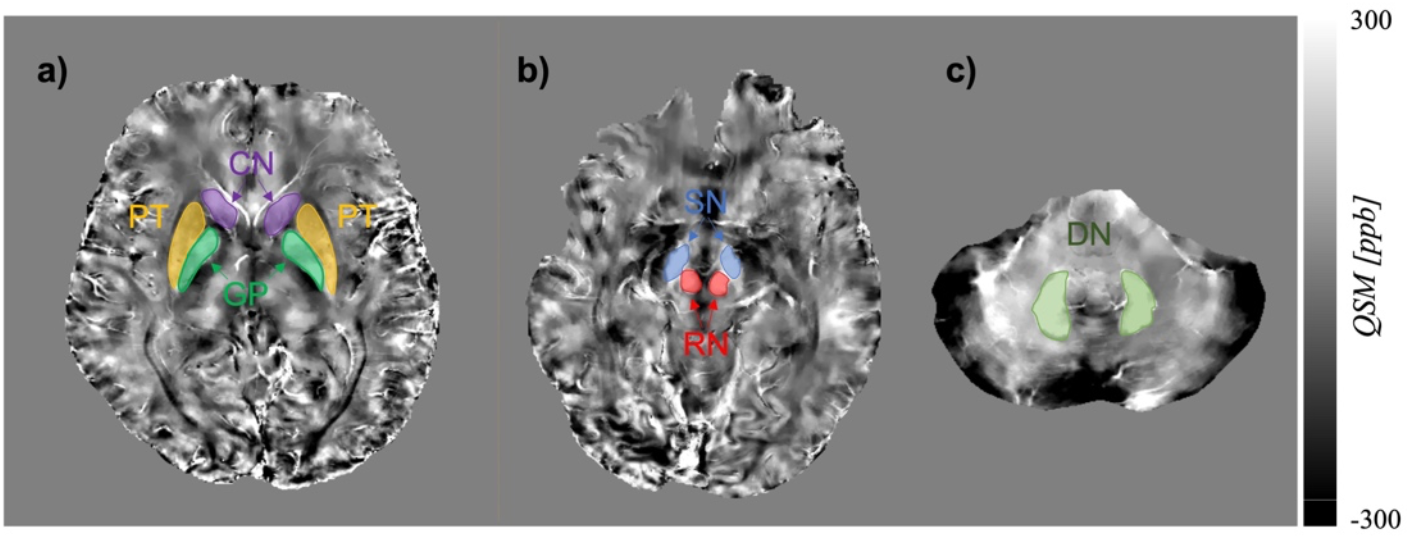
Representative axial QSM images showing manually segmented ROIs used for quantitative analysis of brain iron level. (a) Putamen (PT, yellow), globus pallidus (GP, green), and caudate nucleus (CN, purple); (b) red nucleus (RN, red) and substantia nigra (SN, blue); (c) dentate nucleus (DN, light green). ROIs were segmented based on anatomical reference images and susceptibility values were quantified relative to CSF.

Given that the brains of older participants may exhibit structural differences, such as atrophy, increased air-tissue interface gaps, and overall volume loss compared to the younger individuals used during protocol optimization, we selected three different optimized QSM pipelines for evaluation in the AD and age-matched HC cohorts. These pipelines were constructed using the top performing algorithms from each reconstruction step including phase unwrapping, background field removal, and dipole inversion. The selected pipelines are: (1) Laplacian phase unwrapping, RESHARP background field removal, and MEDI dipole inversion; (2) SEGUE phase unwrapping, RESHARP background field removal, and MEDI dipole inversion; and (3) Laplacian phase unwrapping, VSHARP background field removal, and Star-QSM dipole inversion.

## RESULTS

### Optimized QSM Pipeline

Figure 3 displays susceptibility maps generated from 32 QSM processing pipelines, varying in phase unwrapping, background field removal, and dipole inversion techniques. Overall, visual differences between the Laplacian and SEGUE unwrapping methods were minimal, with the exception of reduced signal intensity near air-tissue interfaces when using SEGUE. This is because SEGUE provides a phase reliability mask that can be used to refine the brain mask, whereas Laplacian-based unwrapping does not offer this capability. Non-linear echo phase combination with Laplacian reduced blooming artifacts, although direct evaluation of unwrapped phase images was not feasible. Background field removal had a significant impact on image quality. Among the four tested methods, VSHARP and RESHARP yielded better results, with RESHARP producing slightly more homogeneity at the cost of some contrast suppression and increased brain boundary erosion. In contrast, LBV and PDF resulted in prominent shadowing artifacts due to residual background fields. Across inversion methods, MEDI consistently produced smooth and visually coherent maps, though some boundary blurring and global smoothing were evident. The iLSQR and TKD maps showed elevated susceptibility values along tissue boundaries, likely due to limitations in managing sharp phase transitions and background field artifacts near anatomical edges. Star-QSM, despite resembling TKD in performance, delivered the best cortical detail and required the least processing time.

**Figure 3.**
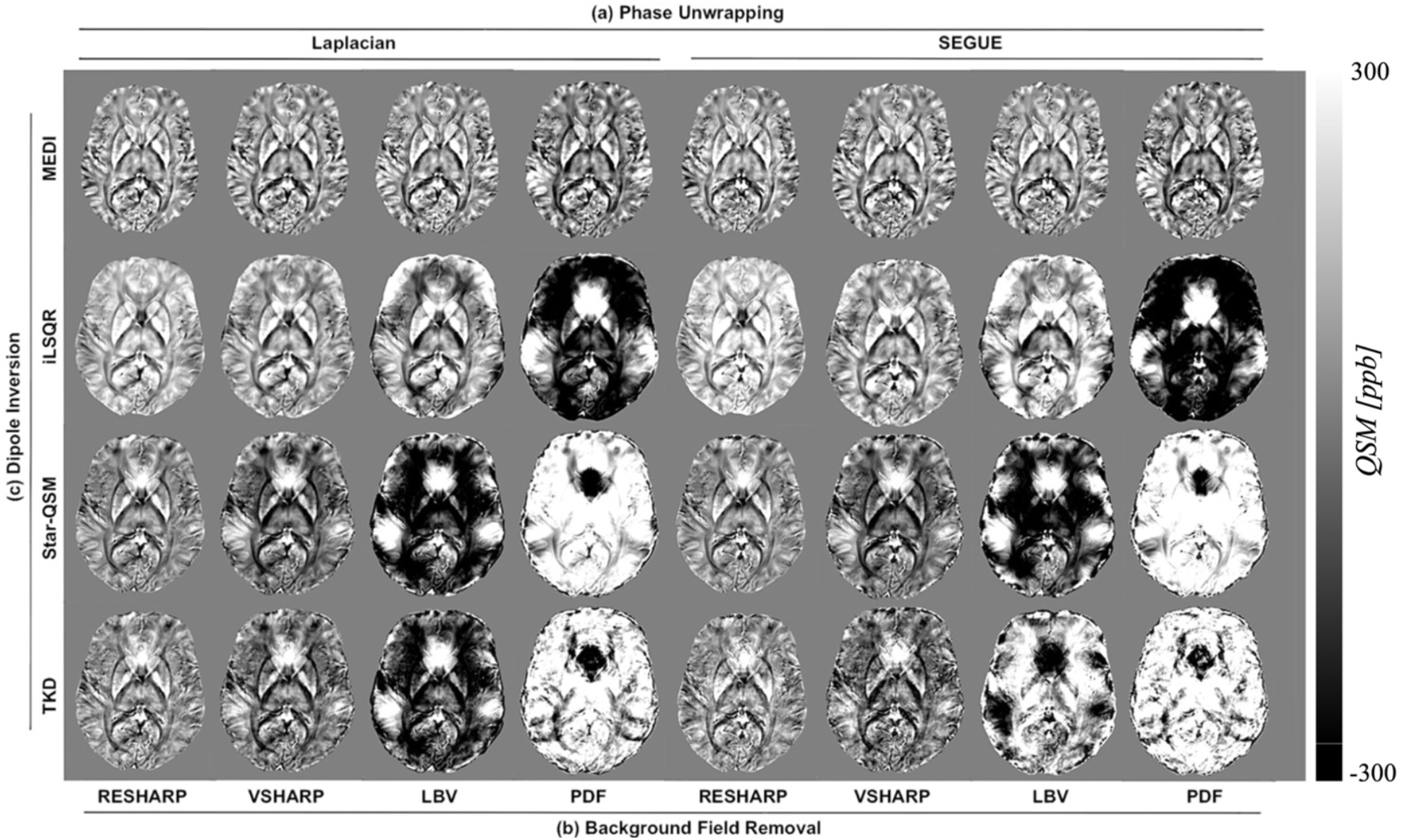
Susceptibility maps generated using 32 different QSM processing pipelines combining two phase unwrapping methods (Laplacian and SEGUE), four background field removal techniques (RESHARP, VSHARP, LBV, and PDF), and four dipole inversion algorithms (MEDI, iLSQR, Star-QSM, and TKD). Each row corresponds to a different dipole inversion method, while columns are grouped by phase unwrapping and background field removal approaches.

Figure 4 presents susceptibility maps acquired at five different in-plane resolutions, focusing on the midbrain region, with zoomed-in views of the CN, PT, GP, and sulci/gyri. Data processing was performed using SEGUE for phase unwrapping, V-SHARP for background field removal, and MEDI for dipole inversion. The first row shows whole-brain QSM images from the same participant at varying resolutions. Box 1 highlights subcortical structures (CN, PT, GP), while Box 2 includes cortical gray and white matter regions. Higher in-plane resolutions provided improved delineation of anatomical boundaries, particularly in PT, GP, and cortical structures. The second and third rows show magnified views of Box 1 and Box 2, respectively. As expected, scanning at mm in-plane resolution improved structural visualization but also resulted in a smaller brain mask. This was primarily due to reduced SNR and increased image non-uniformity, particularly near air-tissue interfaces such as the nasal cavity. These factors contributed to erosion of structures like CN, even when using a permissive brain mask threshold (f = 0.1). While we applied the same brain masking pipeline across all resolutions for consistency, the finer voxel size at 0.2 mm may have led to more conservative masking. Although isotropic resolution is generally recommended for QSM processing, we selected a 0.3 mm in-plane resolution to preserve whole-brain coverage within each axial slice while maintaining structural and quantitative integrity. This approach enables clearer depiction of small anatomical features, supports accurate measurement and quantification, and offers a scan time comparable to standard clinical protocols.

**Figure 4.**
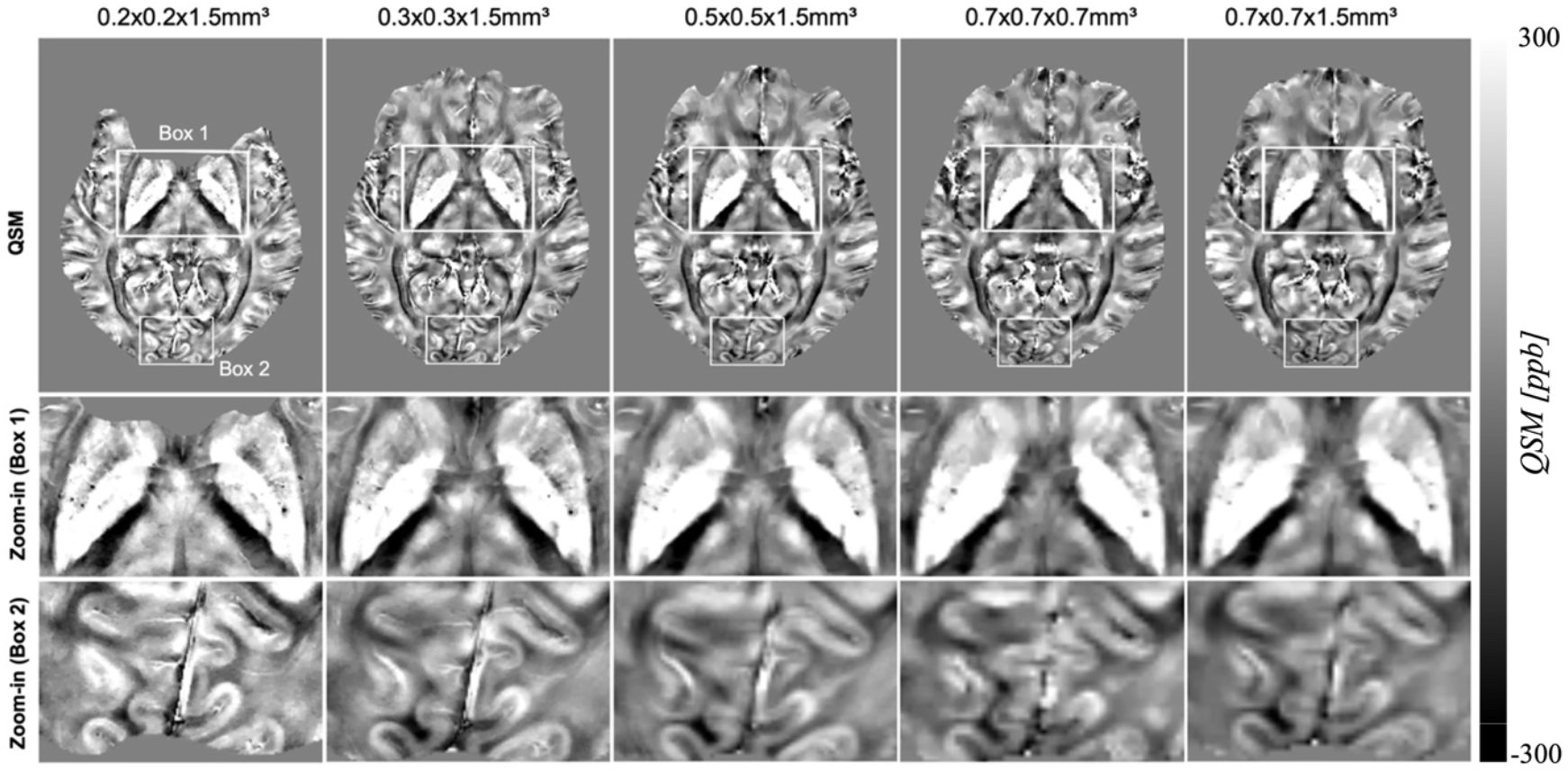
Comparison of QSM maps acquired at five different in-plane resolutions: 0.2×0.2×1.5 mm^3^, 0.3×0.3×1.5 mm^3^, 0.5×0.5×1.5 mm^3^, 0.7×0.7×0.7 mm^3^, and 0.7×0.7×1.5 mm^3^. The top row shows axial QSM slices through the midbrain, with Box 1 highlighting subcortical structures and Box 2 focusing on cortical gray and white matter. The middle and bottom rows display zoomed-in views of Box 1 and Box 2, respectively.

To evaluate the effect of referencing on susceptibility quantification, we processed QSM maps using two approaches: CSF referencing and whole-brain mask referencing. For scans acquired at mm in-plane resolution, both referencing methods produced nearly identical susceptibility maps, with minimal visual or quantitative differences. Given that CSF referencing is more commonly used in the literature, we adopted it for the remainder of the study. However, at 0.2 mm in-plane resolution, CSF referencing often resulted in incomplete or empty susceptibility maps due to signal loss or noise within the ventricles. In such cases, whole-brain or larger-region referencing proved more reliable and was preferred for maintaining map integrity. To assess the influence of VRT on QSM reconstruction quality, we evaluated susceptibility maps generated using three different thresholds: 0.3, 0.5, and 0.7. As VRT increased, more voxels were included in the brain mask, resulting in progressively less erosion of brain tissue and better preservation of peripheral structures (Supplementary Figure 1).

### Iron Assessment in AD Brain

The three pipelines were selected based on quantitative evaluation of the full set of 32 pipelines. Specifically, we assessed each pipeline using metrics such as regional susceptibility consistency, standard deviation within anatomically defined ROIs, artifact suppression, and agreement with known anatomical susceptibility distributions. The three selected pipelines consistently outperformed others across these metrics and were therefore chosen for analysis of iron aggregation on the AD and age-matched HC cohorts. Figure 5 illustrates the results of the three selected QSM processing pipelines, showing axial views of the unwrapped phase, local field maps, and susceptibility maps. All pipelines demonstrated successful phase unwrapping with no significant residual wraps. Notably, Pipeline 2, which used SEGUE for phase unwrapping, produced more homogeneous phase images compared to the Laplacian-based approaches in Pipelines 1 and 3, where a visible brightness gradient was present. Despite these differences, the choice of unwrapping algorithm had minimal effect on background field removal, as all pipelines yielded comparable local field maps. However, VSHARP in Pipeline 3 revealed more distinct hyper- and hypointensities when paired with SEGUE than with Laplacian, suggesting a sensitivity to the phase unwrapping method. Regarding dipole inversion, Star-QSM (Pipeline 3) demonstrated superior capability in resolving hypointensities compared to MEDI (used in Pipelines 1 and 2). Star-QSM performance remained consistent with AD and HC, particularly showing expected erosion in the DN region. Nonetheless, hyperintensities observed in the RN and SN regions posed challenges for resolution. Overall, the combination of Laplacian unwrapping, VSHARP background removal, and Star-QSM (Pipeline 3) showed improved visualization, though MEDI maintained robust and reliable performance across subjects.

**Figure 5.**
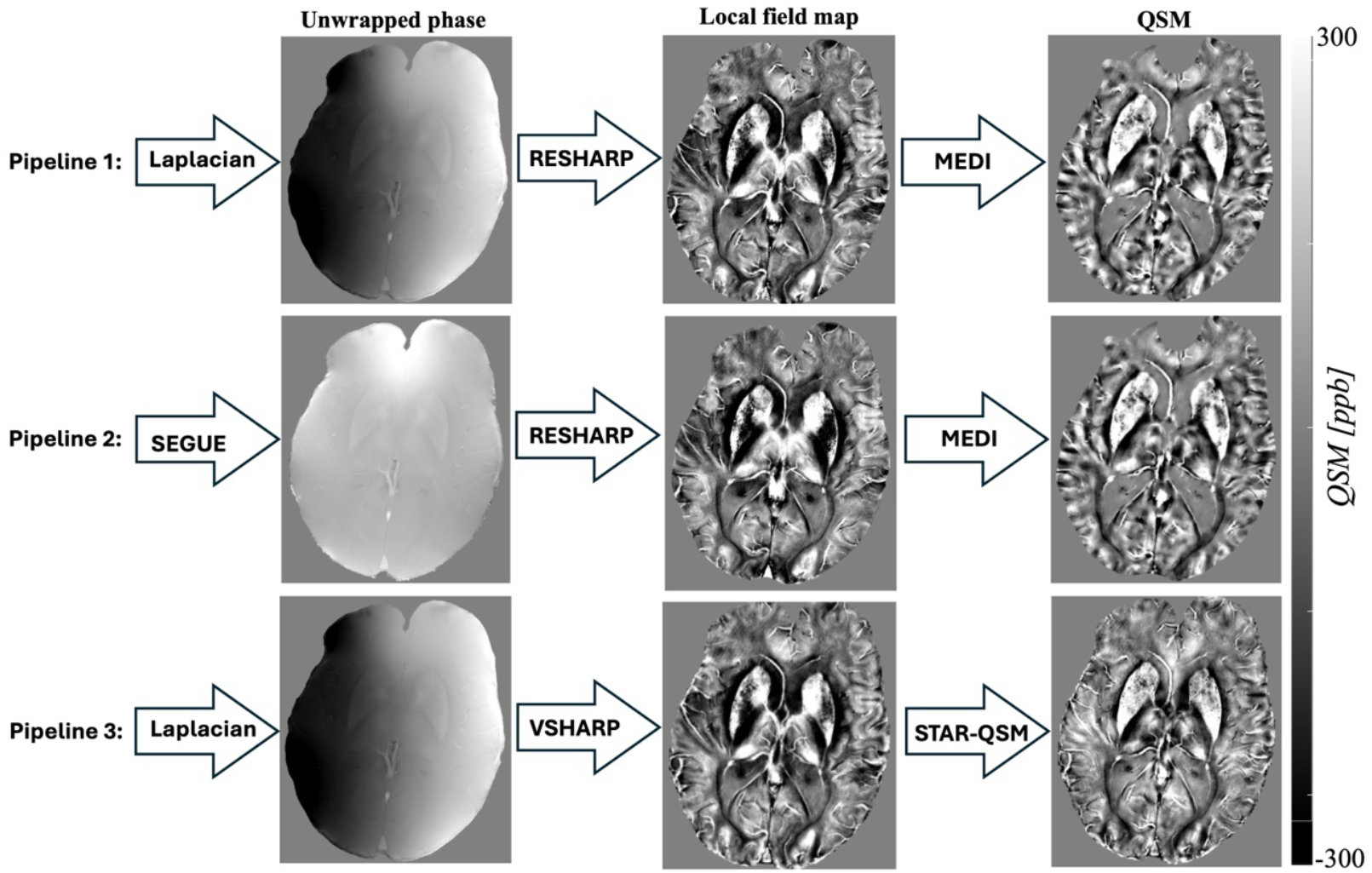
Comparison of three QSM processing pipelines applied to the AD and age-matched HC cohorts, showing axial slices of unwrapped phase images, local field maps, and susceptibility maps. Differences emerged at the QSM stage, with STAR-QSM better resolving hypointensities and showing consistent performance.

Quantitative susceptibility values were compared across three QSM pipelines in six deep gray matter ROIs for both AD and healthy HC groups (see Table 2). Overall, Pipeline 3 consistently demonstrated the highest contrast between AD and HC groups and the lowest standard deviations across most ROIs, indicating improved sensitivity and reproducibility. Figure 6 presents representative QSM images processed using pipeline 3, comparing a HC (a) and an individual with AD (b) across three axial brain slices. Key regions of interest including the RN, SN, CN, GP, PT, and DN, are clearly visualized. In the AD case, enhanced susceptibility contrast is evident in the GP and DN regions compared to the healthy subject. Quantitative analysis (using pipeline 3) confirms this visual observation, showing that the AD group exhibits elevated susceptibility values in the GP (227.5 ppb) and DN (221.7 ppb) relative to HCs (124.6 ppb for GP and 168.9 ppb for DN). Differences were observed in susceptibility values among the other ROIs reported in Table 2. These findings suggest increased iron accumulation in specific deep brain structures in AD, potentially serving as biomarkers for neurodegeneration.

**Table 2:** Susceptibility, χ (ppb) of various anatomical structures relative to CSF using three different pipelines.

| Anatomical Structure | Pipeline 1 |  | Pipeline 2 |  | Pipeline 3 |  |
| --- | --- | --- | --- | --- | --- | --- |
|  | AD | HC | AD | HC | AD | HC |
| Caudate nucleus (CN) | 88.4 ± 8.2 | 85.1 ± 7.7 | 89.2 ± 7.5 | 84.7 ± 6.9 | 91.3 ± 5.9 | 85.2 ± 5.1 |
| Putamen (PT) | 87.3 ± 7.1 | 83.5 ± 6.5 | 88.6 ± 6.9 | 83.9 ± 5.9 | 89.6 ± 6.8 | 84.3 ± 5.4 |
| Globus pallidus (GP) | 219.1 ± 14.6 | 120.4 ± 11.3 | 222.8 ± 13.2 | 122.5 ± 10.6 | 227.5 ± 11.9 | 124.6 ± 10.8 |
| Red nucleus (RN) | 139.8 ± 10.3 | 129.7 ± 9.6 | 141.2 ± 9.7 | 130.1 ± 8.7 | 142.3 ± 9.1 | 130.3 ± 8.5 |
| Substantia nigra (SN) | 165.3 ± 8.1 | 153.9 ± 7.2 | 167.2 ± 7.7 | 155.4 ± 6.8 | 168.4 ± 7.5 | 156.4 ± 6.7 |
| Dentate nucleus (DN) | 216.4 ± 13.8 | 165.3 ± 13.2 | 219.7 ± 12.6 | 167.4 ± 11.8 | 221.7 ± 11.2 | 168.9 ± 11.3 |

**Figure 6.**
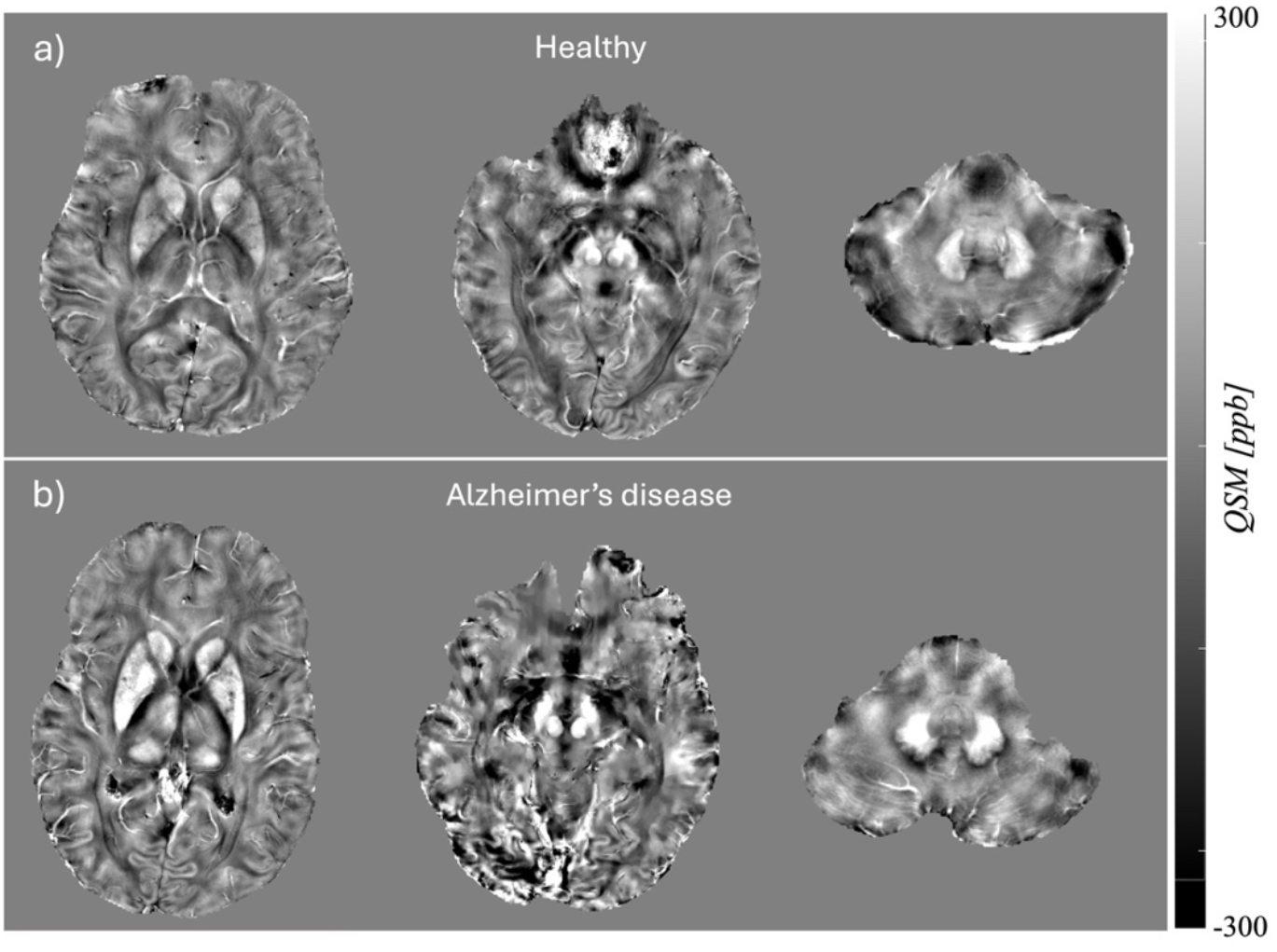
Representative QSM images from (a) a HC and (b) an individual with AD, shown across three axial brain slices. Increased susceptibility is visually apparent in the GP and DN of the AD subject compared to the HC. Quantitative analysis confirmed these observations, with the AD group showing elevated mean susceptibility values in the GP and DN, indicative of greater iron accumulation in these regions.

## DISCUSSION

This study presents an optimized QSM processing pipeline tailored for 7T MRI and demonstrates its feasibility and performance in detecting susceptibility changes in individuals with AD compared to age-matched HCs. Given the increased prevalence of iron accumulation in the aging brain and its association with neurodegeneration, the accurate and reliable mapping of magnetic susceptibility using QSM is essential for advancing biomarker discovery in AD.

In this study, we followed the implementation recommendations outlined in the consensus guidelines published by the ISMRM Electro-Magnetic Tissue Properties Study Group on QSM for clinical brain research [46]. As our study focused on UHF MRI systems, we evaluated and tested multiple QSM postprocessing pipelines to identify an optimized approach suitable for the increased signal sensitivity and specific challenges associated with UHF imaging. While UHF systems offer higher susceptibility sensitivity, they also introduce complexities such as B_0_ and B_1_ field nonuniformities, which can result in signal loss near tissue-air interfaces and nonuniform image contrast [40]. These factors underscore the importance of adapting standard QSM protocols and highlight the need for dedicated QSM implementation guidelines tailored specifically for UHF MRI.

We evaluated 32 QSM pipelines and systematically compared different algorithmic combinations for phase unwrapping, background field removal, and dipole inversion. Among these, three top-performing pipelines were selected based on quantitative metrics and visual quality, and anatomical fidelity. To perform our analysis, we utilized SEPIA, a pipeline processing tool specifically designed to integrate a wide variety of QSM tools within a single platform. While it provides core QSM methods for essential processing, SEPIA primarily supports external tools and implementations developed by the broader QSM research community. This integration allowed us to efficiently compare and evaluate multiple pipeline combinations, ensuring consistency in data handling and reproducibility across different processing approaches. Our findings highlight the significance of choosing a well-matched set of reconstruction steps, as performance varied depending on algorithm compatibility.

In the phase unwrapping step, SEGUE produced smoother and more homogeneous phase images but showed slightly reduced signal intensity near air-tissue interfaces, whereas Laplacian-based unwrapping preserved more signal integrity in such regions. Among background field removal methods, RESHARP demonstrated improved image uniformity, while VSHARP delivered higher contrast at the expense of some variability in susceptibility measurements when paired with different unwrapping techniques. In terms of inversion techniques, MEDI yielded smoother and anatomically coherent maps and was especially robust in resolving hypointensities in deep gray matter regions. Star-QSM, on the other hand, provided better cortical detail and faster computation, making it suitable for time-sensitive applications. These findings support the notion that pipeline performance is contingent on algorithmic synergy and should be carefully tuned, particularly when imaging populations with age-related atrophy, enlarged CSF spaces, and increased air-tissue interfaces.

The optimized QSM pipeline (Laplacian + SHARP + STAR-QSM) was tested in AD and aged-matched HC populations. Quantitative analysis revealed significantly higher susceptibility values in the GP and DN and slightly higher susceptibility values in the other deep gray matter nuclei in the AD group compared to HCs, suggesting elevated iron accumulation in these regions. These findings are consistent with previous studies, including a 3T MRI study that reported regional associations between magnetic susceptibility and aging, cognitive function, and PET markers of amyloid and tau in the basal ganglia and inferior gray nuclei [32, 59]. However, consistent susceptibility changes in cortical regions were not observed. This supports our current observation that deep gray matter structures exhibit greater sensitivity to QSM-detectable changes compared to cortical areas. In addition, another study showed that QSM at 3T can detect brain iron accumulation patterns across AD progression and suggested that these changes may reflect secondary neuronal damage from oxidative stress [11]. Notably, they also reported QSM changes in cerebellar regions, an area where further investigation is warranted. Further supporting the clinical value of QSM, it was shown that magnetic susceptibility was more sensitive than gray matter volume in differentiating between cognitively normal, amnestic MCI, and AD individuals [60]. Importantly, QSM was especially effective in distinguishing cognitively normal from amnestic MCI, a distinction that gray matter volume could not as reliably achieve. This suggests that susceptibility changes due to iron accumulation may precede gross volumetric changes, highlighting the potential of QSM as an early biomarker. Finally, recent advances such as the harmonized UK7T protocol and pipeline have demonstrated a threefold improvement in reproducibility of QSM and R2* at 7T compared to 3T [41]. These results emphasize the readiness of ultra-high-field protocols for clinical and multi-site applications, reinforcing the clinical translatability of our 7T QSM framework. Our study, using 7T QSM, reinforces the sensitivity of high-field imaging for detecting such nuanced regional variations, and future analyses may focus more deeply on cerebellar structures.

Our results on optimizing 7T QSM pipeline provides a reproducible and robust framework for studying brain iron accumulation in aging and AD. It is well-aligned with current multi-site reproducibility efforts and leverages the enhanced spatial resolution and SNR of 7T MRI to capture subtle pathophysiological changes. The integration of our findings with prior research highlights the growing potential of QSM as a potential tool for diagnosis, disease monitoring, and guiding therapeutic interventions in AD.

## CONCLUSION

In this study, we developed and validated an optimized QSM reconstruction pipeline for 7T MRI and applied it to assess iron accumulation in patients with AD. Through qualitative and quantitative analysis, we demonstrated that specific combinations of unwrapping, background field removal, and dipole inversion methods, particularly Laplacian unwrapping, VSHARP removal, and STAR-QSM inversion yield better susceptibility maps that are sensitive to iron-related changes in AD. This pipeline demonstrated better performance in differentiating AD from HC based on both quantitative fidelity metrics and group-level statistical comparisons. Specifically, it yielded higher contrast and more consistent susceptibility values across anatomically defined regions known to be affected in AD. Our results highlight the importance of tailoring QSM workflows to specific research and clinical goals, and show that 7T QSM offers valuable insights into the iron-related pathology of neurodegenerative disease. By comparing our findings to existing literature, including hybrid PET/MRI and 7T imaging studies, we confirm the relevance of susceptibility-based biomarkers in AD and support the integration of QSM into multimodal imaging frameworks. Future work should focus on validating these findings in larger cohorts, exploring the longitudinal progression of susceptibility changes, and integrating QSM with molecular imaging markers to improve diagnostic precision and treatment monitoring in AD.

## ACKNOWLEDEMENT

The authors would like to thank research coordinators Sarah Binder and Aislinn Diaz helping with the recruitment. This study was supported by a Developmental Project award from Mount Sinai ADRC (P30 AG066514) NIA/NIH and K01 AG075178-01 NIA/NIH grants.

## Notes

### Competing Interest Statement

The authors have declared no competing interest.

